# Not a global map, but a local hash: grid cells decorrelate the representation of position and scramble long-range distance information

**DOI:** 10.1101/2025.09.19.677454

**Authors:** Rishidev Chaudhuri, Yicong Zheng, Randall C. O’Reilly, Charan Ranganath

## Abstract

Grid cells in the medial entorhinal cortex construct an intriguing multiperiodic representation of space whose properties have been the subject of much theoretical speculation. Here we combine modeling with analyses of neural population data from mice and rats to show that the grid cell representation is ideally set up to decorrelate and assign easily distinguishable labels to inputs, thus acting as a pattern separation device, much like a hash function in computing rather than a global map or metric. The multiple modules of the grid cell system allow the threshold for pattern separation to be controlled. We also extend these arguments to show how grid cells can perform pattern separation in abstract and higher-dimensional spaces. This pattern separation ability may serve to enhance episodic memory in the hippocampal formation by reducing interference between similar patterns.

## 1 Introduction

Grid cells in the medial entorhinal cortex integrate self-motion and environmental cues to maintain an ongoing representation of an animal’s spatial location^1–5^. They form this representation using an unusual modular periodic code. Each grid cell encodes location periodically, firing at the vertices of a hexagonal lattice. Grid cells are grouped into a small number of modules—each cell within a module shares the same spatial period but is active at a different phase or offset of the lattice^6^. The superficial layers of the entorhinal cortex, in which the bulk of grid cells are found, provide the dominant input to the hippocampus^7^, a structure implicated in both episodic memory and the representation of space. The unusual properties of the multiperiodic grid cell population code and its location in the gateway to the hippocampus have led to extensive theoretical speculation as to its function^2;8^.

One prominent set of interpretations of grid cell function relies on the similarity between the lattice-like representations and a coordinate grid, suggesting that grid cells provide a global coordinate system and metric for space, in service of a global “map” ^1;9–15^. In this view, the coordinate system and metric allow not just the representation of position and local relationships but also computation of global distances between distant points and long-range navigational route planning. However, extracting long-range distances and navigational vectors from the multiperiodic grid cell code requires computationally intricate algorithms or large numbers of readout neurons^11;15–19^. Moreover, the lattice structure of grid cells can be highly distorted by environmental features such as boundaries and rewards, and is dynamic, showing experience and history dependent^6;20–26^. Thus, the extent to which grid cells play a role in global distance measurements and navigational route planning as opposed to other spatial computations is debated.

A second set of ideas for grid cell function focuses on questions of memory capacity and errorcorrection, motivated in part by the hippocampal formation’s well-established role in episodic memory. These ideas highlight the grid cell population code’s remarkable ability to efficiently encode and robustly retrieve a very large set of states, as a result of its modular structure and attractor dynamics^8;11;27–33^. A natural (though sometimes underappreciated) correlate of this high capacity is that the grid cell population representation scrambles or decorrelates long-range metric information, so that except for very nearby locations, distances between locations in physical space are very different from the distances between their corresponding representations in grid cell activity^11;29^. More recently, work on high grid cell capacity has been extended to show how grid cells could provide a repertoire of prespecified stable states that may act as a scaffold for memory representations in the broader hippocampal formation^34^. Instead of a coordinate system, then, these studies suggest the computational metaphor of a hash function—an object that assigns easily separable labels to a large number of input states and, in doing so, scrambles the similarity structure of its inputs.

A possible synthesis between these two theoretical perspectives is provided by observations that grid cells simultaneously preserve very local distance information (on the scale of the width of single neuron firing fields) while scrambling distance information on longer distance scales (greater than the width of single neuron firing fields). This unusual combination of properties could allow grid cells to both provide a set of local (but not global) metrics and maps^26^ and show very high representational capacity^8;11;27–33^, acting much like variants of hash functions called locality-sensitive hashes and locality-preserving hashes^34^. This high-capacity local, rather than global, map picture fits well with the episodic memory function of the hippocampus—encoding specific details associated with individual episodes rather than encoding global relationships between all episodes with respect to each other.

In this study, we empirically test the idea that grid cells act to scramble and decorrelate relationships between input patterns, with the exception of very local structure. We first use the notion of a population vector *similarity kernel* to summarize and formalize previous theoretical observations of decorrelation by grid cells^11;29^ (Section 2.1). We then show that the threshold for decorrelation of spatial locations should vary systematically as a function of anatomical location, thus providing both fine and coarse scale decorrelation and a possible mechanism to control the tradeoff between local structure preservation and long-range pattern separation (Section 2.2). We next use electrophysiological recordings from rats and mice to empirically estimate the similarity kernel and confirm that the grid cell system indeed scrambles distance information, except on very local scales, in accordance with theoretical predictions (Section 2.3). Finally, we extend these ideas to grid cells in higher-dimensional and abstract spaces (Section 2.4).

## 2 Results

### 2.1 The grid cell representation acts to scramble distance relationships while providing well-separated linearly-distinguishable labels to spatial locations

Prior work on the grid cell system has highlighted its ability to robustly encode a very large number of spatial states^11;27–31;34^. As previously argued, a natural consequence of such high capacity and robustness is the decorrelation and interleaving of spatial representations—nearby spatial states can have quite dissimilar representations in grid cell activity, and distant spatial states can have comparatively similar representations in grid cell activity (particularly at the level of a single module)^11;29^.

In this section, we frame such decorrelation in terms of a function, *k*, called a similarity kernel. Given a pair of spatial locations, say ***x***_**1**_ and ***x***_**2**_, *k*(***x***_**1**_, ***x***_**2**_) measures how similar the grid cell network’s encodings of these locations are to each other. Such kernels have been widely and fruitfully used to measure and compare the similarity structure of population representations, often under the name Representational Similarity Analysis (RSA)^35^, as well as to understand which particular features of the world population representations in different brain regions are optimized for and what inductive biases these representations bring to learning ^36^.

Decorrelation implies that the representation by grid cell activity acts to scramble distances, so that the value of the similarity kernel *k*(***x***_**1**_, ***x***_**2**_) is nearly uncorrelated with the physical distance between ***x***_**1**_ and ***x***_**2**_. In particular, beyond a certain distance threshold, nearby points are mapped to well-separated, near-orthogonal population vectors in grid cell space, so that *k*(***x***_**1**_, ***x***_**2**_) takes a nearzero value. We note that these arguments imply that distance relationships are not just scrambled but *homogenized* —the similarity of the representation in grid cell space is close to a small constant value independent of physical distance and the values of ***x***_**1**_ and ***x***_**2**_. These conclusions are shown in Fig. 1 using simulated grid cell population data. The remainder of the section presents a theoretical argument in support of this conclusion.

**Figure 1.**
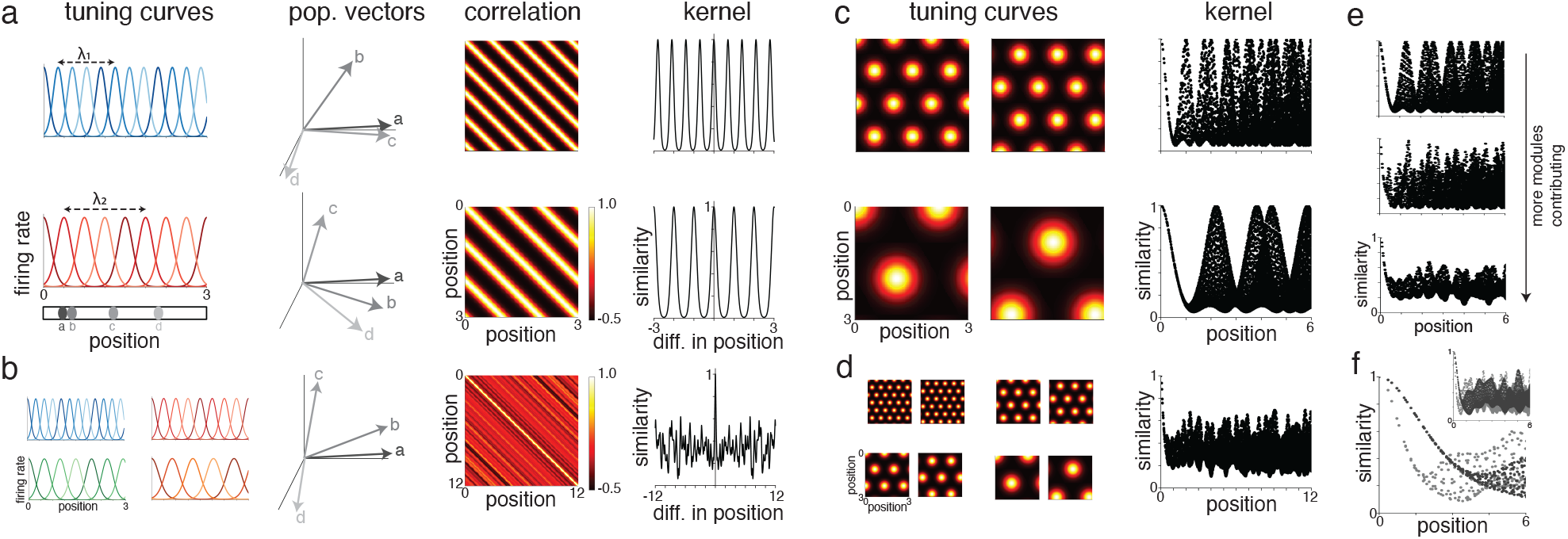
Simulated grid cells produce decorrelated representations of input patterns. (a) First column: tuning curves from two simulated 1D grid cell modules, one with a smaller period (top, in blue) and the other with a larger period (bottom, in green). Each Gaussian-shaped trace shows one neuron. The positions indicated below in gray (labeled *a*-*d*) show the locations corresponding to the population vectors in the second column. Treating position *a* as the reference, position *b* is very close to position *a*, position *c* is close (but not immediately adjacent) to position *a*, and position *d* is distant from position *a*. Second column: schematics of population activity vector in each module corresponding to each of the four locations shown in first column. The population vector for position *b* is correlated with the population vector for position *a*, with the degree of correlation depending on the width of tuning curves in that module. For most modules, the population vectors for position *c* and *d* are near orthogonal to the population vector for position *a* but, because of the periodicity, for some modules these vectors are correlated (e.g. position *c* for module 1). Third column: correlation of population activity vectors for pairs of positions for the two modules. Fourth column: similarity kernel for pairs of points as a function of the distance between them. (b) Measures as in panel (a) but for population activity across all modules. Tuning curves from a subset of modules are shown in first column, population vector schematic in second column, correlation between population activity vectors in third column, and similarity kernel in the fourth. When aggregated across modules, the population activity vectors for *a* and *b* are correlated while the population vectors for *a* and *c, d* are near-orthogonal. Correspondingly, the kernel is sharply localized around 0. (c) Results for simulated 2D grid cell modules. Top and bottom show two modules with different periods. First column: tuning curves of two sample neurons in each module. Second column: similarity kernel against distance for pairs of points. (d) As in (c) but combined across modules. (e) Top to bottom show similarity kernels resulting from combining increasing numbers of modules. As more modules are combined, kernel becomes increasingly localized around 0 and the peaks away from 0 vanish. (f) Similarity kernel when only considering modules with shorter periods (light gray) or larger periods (dark grey). The corresponding kernel shows a faster or slower decay around 0. Inset shows kernel over an extended range.

The argument showing this scrambling is a recapitulation of previous work on high capacity representations in the grid cell system^11;29^, with a focus on estimating the structure of the similarity kernel *k*. To present the argument in its essential form, we make a set of simplifying assumptions. First, we assume that the grid cell population activity is entirely determined by spatial location in a given context (rather than say, the presence of boundaries, velocity, trajectory history, and so on). Thus, if the subject is at spatial location ***x***, the grid cell network’s population activity can be expressed as ***g***(***x***), for some function ***g***^a^. Second, we consider a simplified one-dimensional model of grid cell population activity. Note that results generalize straightforwardly to two dimensions, as we show using simulated data in Fig. 1c,d and using neural population recordings in Fig. 2. In Section 2.4 we moreover extend the argument to putative higher-dimensional grid cells and show that the same conclusions hold. For now, however, we assume that the grid cell population encodes a 1-D dimensional spatial variable, *x*.

**Figure 2.**
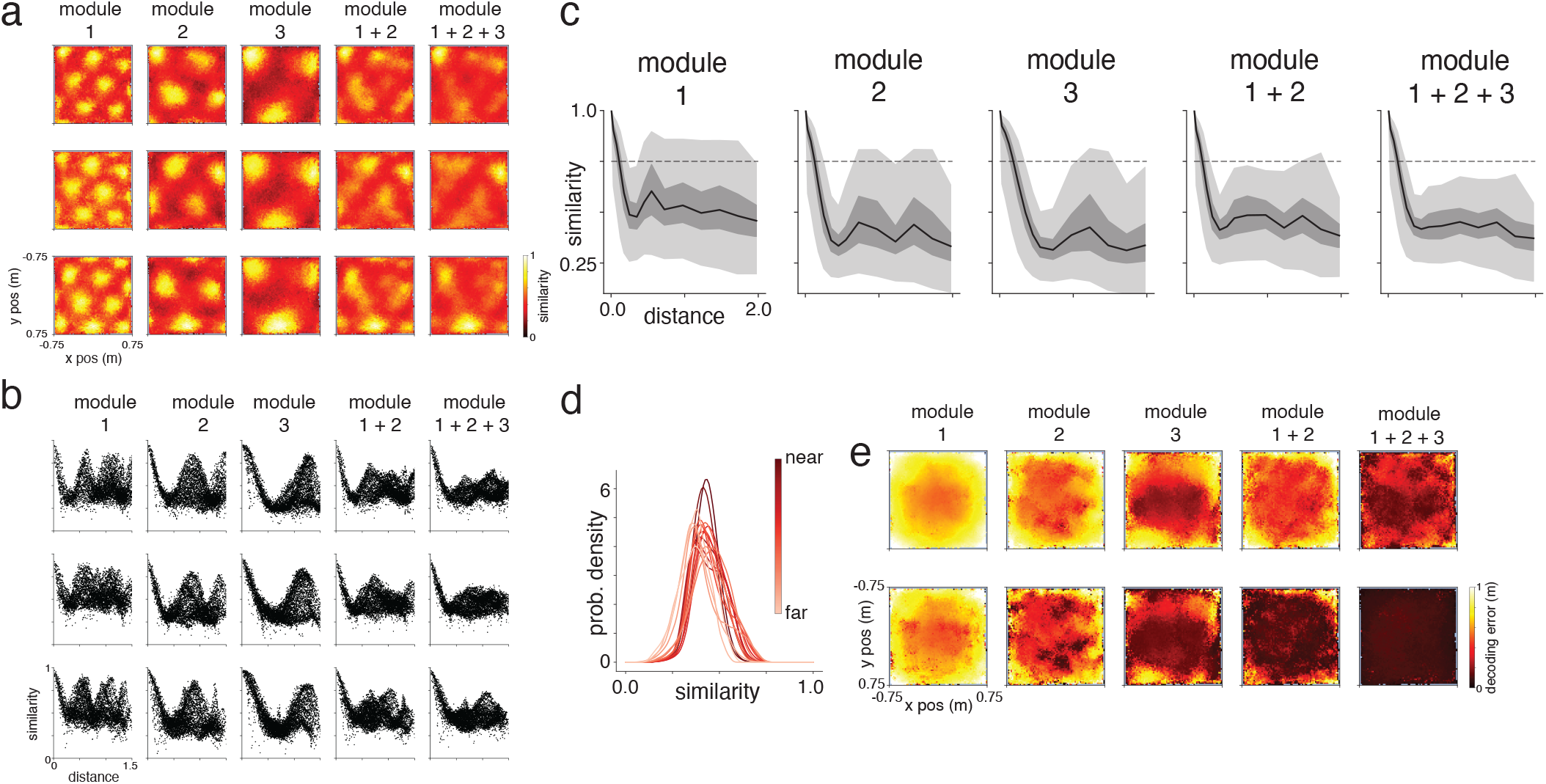
Similarity structure of grid cell representation from study of Gardner et al. (2022). Panels (a) and (b) show results for kernels from particular locations, while (c)-(e) show aggregate statistics across locations. (a) Each row shows similarity kernel (i.e., dot product) of the grid cell representation between a specific fixed location (indicated as blue star in final column) and all other locations in the arena. Each column shows a different module or combination of modules: first three columns show the three modules from the study, fourth column shows combination of modules 1 and 2, and final column shows combination of all three modules. Thus, for example, the heatmap on the first row and second column is 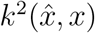, where 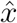 is a particular location in the arena and *k*^2^ is the kernel induced by the second module. Kernels become increasingly localized as modules are combined (i.e., moving left to right). (b) Each row shows similarity kernel of the grid cell representation between a specific location (same locations as in (a)) and all other locations in the arena, plotted as a function of their physical distance from each other. Columns as in (a). As modules are combined, except for very nearby points, grid cell similarity is increasingly independent of physical distance and concentrated about its mean value. (c) Similarity kernel for all pairs of locations as a function of distance. Black line shows median of the distribution, dark grey region is 25th to 75th percentile, and light grey region is 1st to 99th percentile. (d) Histograms of grid cell similarity for all three modules combined, for all pairs of points that are not very close (i.e., beyond the threshold for decorrelation). Each trace shows the histogram for a certain distance bin, ranging from pairs of points 0.3-0.4m apart (darkest trace) to pairs of point 1.9-2m apart (lightest trace). Distributions are highly overlapping and not structured by distance. (e) Error of a threshold linear decoder that uses the population vector to decode position in the arena, shown for a threshold of 0.65 (top) and 0.75 (bottom).

Let the grid cell population consist of *M* modules, with the spatial scale of the *m*-th module being *λ*_*m*_, and the vector of population activity of the *m*-th module being denoted by the vector ***g***^***m***^(*x*). The set of module periods are {*λ*_1_, …, *λ*_*M*_ }. We assume that each module contains *N* neurons, of which only one is active at any given time (corresponding to the phase of the grid cell representation). This is a simplified representation of the tuning curves. If the spatial location represented is *x*, this representation means that the population activity vector for the *m*-th population has a 1 in the position corresponding to ⌈*N* (*x/λ*_*m*_ mod 1) ⌉, where ⌈·⌉ represents the ceiling function that rounds a decimal up to an integer. Thus, for example, if *λ*_*m*_ is 2 and *N* is 4, spatial locations in [0, 0.5], [2, 2.5], [4, 4.5] and so on correspond to the first neuron in ***g***^***m***^(*x*) being active and the rest being 0; spatial locations in [0.5, 1], [2.5, 3], [4.5, 5] and so on correspond to the second neuron in ***g***^***m***^(*x*) being active; and similarly for the remaining two neurons in the module. In Fig. 1a, we show tuning curves and population vectors for two modules from such a model, though using more realistic Gaussian tuning curve shapes, and in Fig. 1c we show the 2D case.

The activity vector of the grid cell population as whole, ***g***(*x*), is simply the concatenation of the individual ***g***^*m*^ vectors, and thus is a vector of length *M* × *N* (schematic in Fig. 1b, d).

Given a pair of spatial locations *x*_1_ and *x*_2_, define the *similarity kernel k*^*m*^ for module *m* to be the dot product between the corresponding vectors of population activity for neurons in that module. That is,

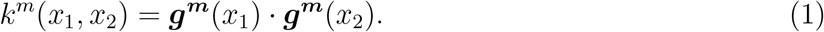

And define the kernel function for the population as a whole to be

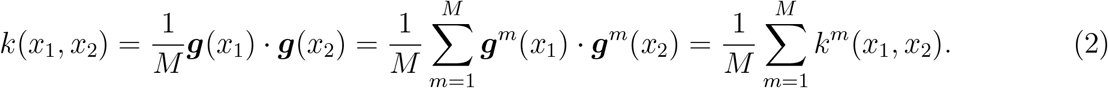

Here the normalization factor of 1*/M* simply serves to remove the dependence on the number of modules.

These kernel functions capture the similarity of the grid cell representations for the positions *x*_1_ and *x*_2_ as measured by their dot product or (non-mean-subtracted) correlation (see examples in Fig. 1a-d right columns). If the grid cell population activity vectors for positions *x*_1_ and *x*_2_ (i.e., ***g***(*x*_1_), ***g***(*x*_2_)) are similar, then the vectors point in similar directions and the dot product is high, yielding a high value for the kernel function. If the population activity vectors are very dissimilar, then they point in different directions leading to a small dot product and hence small value of the kernel.

We next analyze the structure of these kernel functions, first for a single module and then for combinations of modules, again following the structure of the argument used in previous work on high capacity representation in the grid cell system^11;29^.

Note that the kernel function for a single module, *k*^*m*^(*x*_1_, *x*_2_), is non-zero and takes value 1 if the corresponding positions *x*_1_ and *x*_2_ lie at the same phase of the grid cell representation for module *m* and takes value 0 otherwise. We distinguish three possible situations.

First, if *x*_2_ is immediately adjacent to *x*_1_, meaning that they are separated by less than the typical firing field width of a grid cell (≈*λ*_*m*_*/N* in our simplified model), then they will be assigned to the same phase of the grid cell representation (i.e., the same bin). In our simplified model, in this situation ***g***^*m*^(*x*_1_) = ***g***^*m*^(*x*_2_), and thus *k*^*m*^(*x*_1_, *x*_2_) = 1.

Second, if *x*_2_ is close (but not extremely close) to *x*_1_, meaning that it is further than the typical width of a grid cell firing field but within the period of the module (i.e., *λ*_*m*_*/N <* |*x*_2_−*x*_1_|*< λ*_*m*_), then the corresponding population vectors for this module will be non-overlapping and hence orthogonal, with *k*^*m*^(*x*_1_, *x*_2_) = 0.

Finally, consider a random point lying outside the period of the module (|*x*_2_ *x*_1_| ≥ *λ*_*m*_). With probability *p* = 1*/N*, these points will be assigned to the same phase of the grid cell representation for that module yielding *k*^*m*^(*x*_1_, *x*_2_) = 1. With probability 1 −*p*, the positions are at different phases and the population activity does not overlap. Thus with probability 1 − *p* the population vectors are orthogonal and *k*^*m*^(*x*_1_, *x*_2_) = 0.

Note that, even with 1 module, distance relationships are already scrambled—nearby spatial locations map to orthogonal population vectors while distant spatial locations can sometimes map to the same or similar population vectors, right panels of Fig. 1a, c.

Next, note that combining across multiple modules preserves the separation of nearby spatial locations while reducing the probability that distant spatial locations map to similar vectors. Thus, multiple modules together provide decorrelated representations across spatial locations.

To see this, consider multiple modules with different spatial scales. As long as the two points *x*_1_ and *x*_2_ are not immediately adjacent to each other (i.e., provided |*x*_1_−*x*_2_| *>*| *λ*_*m*_*/N*) and except for special choices of spatial scales, for two randomly chosen points *x*_1_ and *x*_2_, the probability that the population activity in module 1 overlaps for *x*_1_ and *x*_2_ is independent of the probability that the population activity in module 2 overlap for *x*_1_ and *x*_2_ (see more detailed arguments in prior work^11;29^). That is, the probability that *k*^1^(*x*_1_, *x*_2_) = 1 is independent of the probability that *k*^2^(*x*_1_, *x*_2_) = 1. A similar argument applies for each pair of modules. The population kernel 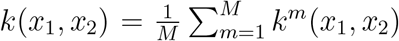 is thus the sum of independent terms that each take value 1 with probability *p* and value 0 with probability 1 − *p*. Consequently, as long as *x* and *x*_2_ are not too close, *k*(*x*_1_, *x*_2_) can be modeled as a binomial random variable, and has mean *p* and standard deviation 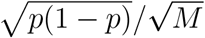.

These results mean that as long as two locations *x*_1_ and *x*_2_ are not extremely close, the population vectors corresponding to these locations will on average be nearly orthogonal to each other, with mean kernel value of *p*, regardless of the distance between *x*_1_ and *x*_2_ (and will be exactly orthogonal to each other for points that are nearby but not extremely close), shown in Fig. 1b, d. Moreover, the dependence of the standard deviation of *k*(*x*_1_, *x*_2_) on *M* implies that as more modules are added the spread of the kernel values around *p* will decrease, and the kernel for two randomly chosen points will be closer and closer to its mean value of *p*, Fig. 1e. Typical fluctuations are on the size of the standard deviation but fluctuations much larger than the standard deviation become exponentially unlikely (as can be shown, e.g., by standard concentration of measure bounds). This concentration means that the grid cell representation acts not just to scramble distance relationships but also to homogenize them to an increasingly narrow band of values around *p*. Thus, the grid cell representation acts to decorrelate population vectors. It moreover does so very efficiently using only a small number of neurons^11;29^.

Given that the similarity kernel *k*(*x*_1_, *x*_2_) measures the dot product between population representations, the small value of *k* for most pairs of points means that the grid cell encodings of *x*_1_ and *x*_2_ are not just well-separated but near-orthogonal. This near-orthogonality means that the representation for each spatial location is linearly separable from the others, allowing downstream neurons to easily use these representations as labels without interference from the labels of other locations, making for efficient readout and memory storage^11;34^.

For example, if ***g*** is the grid cell population vector with *n*th component *g*_*n*_ (note that *n* is the index for neurons not modules), let the activity of a threshold-linear downstream neuron be modeled as,

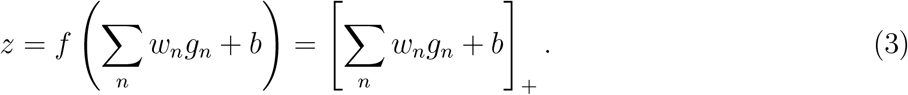

Here *f* is an activation function, which we set to be rectified-linear for simplicity, *w*_*n*_ is the weight of the *n*th grid cell input onto the downstream neuron, and *b* is a bias. To have this neuron be active around a particular location 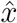 and silent otherwise, one can simply set the weights equal to the (sparse) vector of grid cell population activity corresponding to this location. Thus 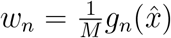, where the normalization factor 1*/M* is included for convenience but is not necessary.

With these weights, the activity of the downstream neuron for any position *x* is a thresholded version of the kernel function and can be written as

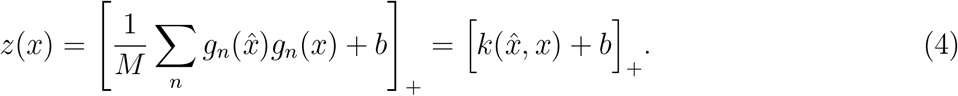

The localization of the kernel function about 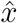 thus means that the activity of the downstream neuron is 0 unless 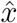 is very close to *x*, with the sharpness of tuning set by the bias parameter *b*. Since the weights correspond to the activity of the presynaptic population at the desired location, they can be learned by a simple correlational learning rule.

We note that this linear separability and the argument in the previous paragraph is a restatement of the well-known idea that grid cell activity could be summed up and thresholded to produce place cell activity^9;11;37;38^, expressed in terms of the similarity kernel function.

The argument so far uses the similarity kernel as measured by the dot product or non-mean-subtracted correlation as a measure of distance between population vectors, but equivalent results hold when distance between grid cell population vectors is measured as squared Euclidean distance, 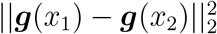 Expressing Euclidean distance in terms of the kernel yields

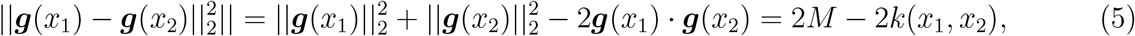

where 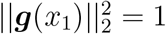because ***g***(*x*_1_) has *M* entries equal to 1 with remaining entries equal to 0 (and similarly for 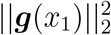). For randomly chosen *x*_1_ and *x*_2_, on average we have

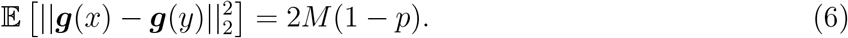

Thus the average distance between population vectors is of a similar magnitude to the length of the population activity vectors themselves (i.e., *M*), meaning that points are very well-separated. Also as before, the standard deviation of distances around this mean value shrinks as the number of modules increases, acting to homogenize distances so that points are mapped to equidistant population vectors. Thus, when measured by Euclidean distance, the grid cell representation serves to embed positions as well separated near-equidistant vectors (and see the discussion for consideration of other similarity measures).

In sum, then, the grid cell representation acts to embed spatial locations in such a way as to assign each location a unique label while scrambling the representation of distances between points that are more than a small distance apart (i.e., the width of a single neuron firing field), and does so efficiently, using a small number of neurons. Thus, once input patterns are more than a certain threshold distance apart, the grid cell representation further separates these patterns. The structure of these representations can be captured by a kernel function *k*, which encodes the similarity of the representation of pairs of spatial locations.

### 2.2 The threshold for decorrelation by grid cells depends on anatomical location

In the previous section, we described the similarity kernel for single grid modules and for the grid cell population as a whole. In particular, the similarity kernel function for the grid cell population as a whole has a certain characteristic width—spatial locations closer than this width have their similarity preserved by the population representation, while spatial locations further apart have their representations decorrelated. In this section, we describe the similarity kernel and its characteristic width for subsets of anatomically contiguous grid cell modules.

The periods and firing field widths of grid cells in rodents vary systematically across modules, ranging from small periods in dorsal medial entorhinal cortex to large periods in ventral medial entorhinal cortex^1;39^. The gradient of scales parallels the increase in place field sizes from dorsal to ventral rodent hippocampus, and it has been suggested that place fields of different size can be formed by preferentially summing subsets of grid cell modules of appropriate size^9;37;38^ (though the mechanistic relationship between grid and place cells is complex). For sub-groups of modules, we here make the similar observation that the gradient of scales creates kernels that in turn have a gradient of widths, moving from similarity kernels that have a low threshold for similarity (and thus aggressively decorrelate or pattern separate inputs) to similarity kernels that have a high threshold for similarity (and thus are more likely to preserve similarity structure and enhance pattern completion by downstream structures).

First, note that each grid cell module performs decorrelation or pattern separation with a different similarity threshold, set by the width of the typical grid cell firing field in that module. In the models from the last section, this width is given by *σλ*_*i*_, where *σ* is the ratio of firing field width to module period. Note that, in accordance with experimental observations, these widths scale with the period of the grid module—grid cells in modules with larger periods have wider firing fields. Spatial locations separated by less than *σλ*_*i*_ are mapped to similar grid cell activity patterns (in the simplified one-hot model described in the previous section they activate the same neuron; more generally they activate overlapping subsets of neurons). Spatial locations separated by more than *σλ*_*i*_ are mapped to orthogonal population vectors with high probability.

Now consider a similarity kernel defined by a weighted sum of module activities. That is,

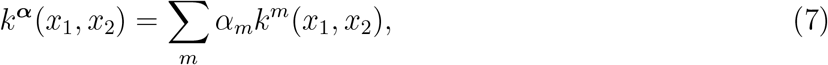

where as before *m* indexes modules, *k*^*m*^(*x*_1_, *x*_2_) is the similarity kernel for the *m*th module, and *α*_*m*_ is the weight for the *m*th module. Note that setting *α*_*m*_ = 1*/M* for all *m* yields the global kernel considered in the previous section. Changing the weights *α*_*m*_ changes the contribution of the kernel from each module and thus allows the similarity kernel to take a variety of shapes; such sums can be used to efficiently approximate a number of translation-invariant functions^40^.

In particular, these weights can control the threshold for decorrelation, Fig. 1f (and see Fig. 5 of Fiete et al. (2008) for a similar observation). Setting weights to be high from dorsal modules with small firing fields overweights sharply decaying kernels and thus leads to pattern separation with a small threshold, so that even slightly dissimilar patterns are decorrelated. On the other hand, setting weights to be high from ventral modules with large firing fields leads to pattern separation with a larger threshold, so that slightly dissimilar patterns are mapped to similar outputs.

As a specific example, consider a downstream neuron modeled as in the previous section as 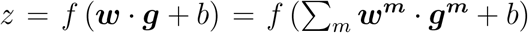, where ***w*** is the vector of weights onto the neuron, ***g*** is the population activity vector of the grid cell population, ***w***^***m***^ is the vector of weights from the *m*-th grid cell module and ***g***^***m***^ is the population activity vector of the m-th module. Note that *m* indexes modules, each of which has *N* neurons. If, as in the previous section, the weights to this downstream neuron are set equal to the activity of the grid cell population at a particular location 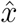, then *z* is a function of the overall population kernel. Instead, let the weights from module *m* be set proportional to the activity of that module at 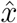, but with weighting factor *α*_*m*_. Thus 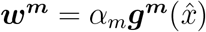. Then,

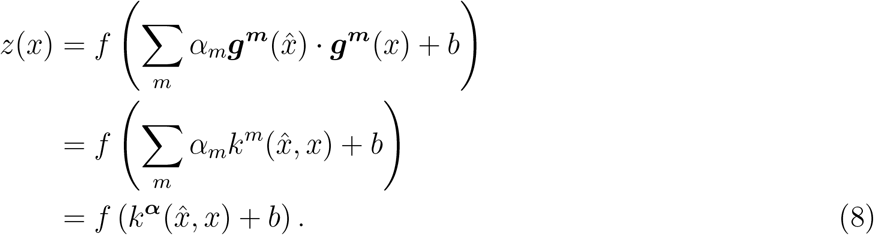

Here *k*^***α***^ is an effective kernel given by the weighted sum of the individual module kernels. As a result, this neuron is maximally active at 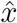 with a response that decays with distance from 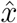 and whose width is determined by the width of the kernel *k*^***α***^. Note that this formulation is equivalent to observations that summing up appropriately sized grid cell modules could yield place cells with different firing field widths, reexpressed in the language of the similarity kernel. However, the perspective is more general than considering a single downstream neuron and applies equally well to readout by a downstream network of neurons rather than a localized readout by a single neuron.

### 2.3 Similarity kernel in grid cell recordings

As highlighted by the framework presented above, arguments for decorrelation by the grid cell code^11;29^, make four key predictions.

#### 1. Scrambled distances beyond a small threshold

Grid cells act to scramble distances in physical space. Thus, with the exception of very nearby points (determined by single neuron firing field widths), similarity structure in grid cell space is uncorrelated with distance in physical space. Moreover, the particular mechanism grid cells use to scramble distances should also lead to distances being homogenized, so that similarity structure in grid cell space should be much more tightly concentrated around its average than distances in physical space.

#### 2. Unique linearly separable labels

While the representation of spatial position at the level of a single module is ambiguous, grid cell population vectors for different spatial positions become increasingly orthogonal as more modules are considered, thus providing unique labels to locations. Moreover, these labels are not just distinguishable, but linearly separable, and position can thus be decoded using a simple linear classifier. Note that this is effectively a restatement of well-known ideas that grid cell activity could be summed up and thresholded to produce place cell activity^9;37;38^ (with the added theoretical guarantee that any location could be assigned a place field). However, our focus is on the linear separability of the patterns whether or not they are thresholded to produce a localized spatial representation.

#### 3. Kernel width varies systematically across an anatomical gradient

Decorrelation can be carried out by subsets of modules, and the spatial threshold for what counts as “very nearby” points (i.e., the width of the similarity kernel) varies systematically depending on which grid cell modules are being sampled.

#### 4. Efficiency

The grid cell representation is extremely efficient at performing this decorrelation, so it should be visible from small numbers of neurons.

To verify these predictions we analyzed electrophysiological population recordings from medial entorhinal cortex in two recently published studies, one containing recordings of multiple grid cell modules from rats foraging in open field environments^41^ and the other containing recordings of putative grid cells from mice running on a one-dimensional virtual reality track^42^.

We first show results from the data of Gardner et al. (2022). These data consist of simultaneous recordings of upto 3 grid cell modules while rats forage in open field environments. We used these data to estimate the similarity kernel function for each module. That is, for each module *m* and pairs of locations *x*_1_, *x*_2_ we computed the dot product of the average population activity vectors at these locations yielding *k*^*m*^(*x*_1_, *x*_2_) = ***g***^***m***^(*x*_1_)· ***g***^***m***^(*x*_2_), where ***g***^***m***^(*x*_1_) is the average population activity vector for module *m* in a small spatial bin centered at *x*_1_ (and similarly for *x*_2_). We also computed the kernel for combinations of modules, using the concatenation of appropriate population vectors.

For example, *k*^1,2^(*x*_1_, *x*_2_) is the similarity kernel imposed by the combination of modules 1 and 2. We calculated this by concatenating the population vectors ***g***^**1**^(*x*_1_) and ***g***^**2**^(*x*_1_) and taking the dot product of this concatenated vector with the concatenation of the population vectors ***g***^**1**^(*x*_2_) and ***g***^**2**^(*x*_2_). Note that this way of combining modules simply weights neurons equally; weighting the different modules or neurons differently would lead to a more flexible set of kernels that interpolate between the kernels imposed by each module.

In Fig. 2a, we show kernels for 3 example spatial locations by plotting the similarity of all locations in the arena with each of these 3 example locations. That is, we plot 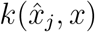 for particular locations 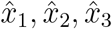, and all locations *x*. At the single module level (first 3 columns), the periodic structure of the grid cell responses is clearly visible and multiple locations have highly correlated population vectors. However, these periods average out when combining across modules, leading to increasingly distinct representations for each location, as in prediction #2 above. Moreover, as can be seen from the increasingly single-peaked and otherwise homogeneous heatmaps, the value of the kernel for pairs of points beyond a small distance threshold approaches a small constant value independent of physical distances, as in prediction #1 above.

In Fig. 2b, we show scatter plots of the value of the kernels from the same 3 example spatial locations as a function of physical distance between locations 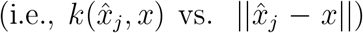, thus directly comparing similarity in grid cell space to distances in physical space. In accordance with prediction #1, except for very nearby points, distances are scrambled even at the level of one module. Moreover, upon combining modules the distances are increasingly homogenous, showing a much smaller spread of values than distances in physical space (i.e., compare the spread of points in the *y* direction for the first 3 columns against the last 2 columns).

Next, to compare similarity in grid cell space to distances in physical space for all pairs of points, we computed the value of the kernel *k*(*x, y*) for all pairs of physical locations *x* and *y*. In Fig. 2c, we plot the distributions of these kernel values as a function of the spatial distance between points *x* and *y*. As with the example locations, for distances beyond the width of a typical firing field, similarity in grid cell space is only weakly correlated with distance in physical space. Moreover, when combined across modules, similarities in grid cell space are concentrated in an increasingly narrow band around the mean value thus verifying that distances are homogenized and not just scrambled, Fig. 2d. Thus, these analyses support prediction #1 above.

Finally, to test the linear separability of locations we trained a simple threshold-linear binary decoder to decode a particular spatial position. Training was carried out using a Hebbian learning rule so that the weights of the decoder for a particular position are set to be the population activity vector at that position. The decoder thus reports that the animal is at that particular position whenever the similarity of the population vector with the weights exceeds a certain threshold. We repeated this procedure for every spatial location. The plots in Fig. 2e show the error of such a decoder at each spatial position in the arena, with the top and bottom rows showing two different decoding thresholds. The decoding error is large for single modules (first 3 columns) and rapidly becomes small as modules are combined (last 2 columns).

Taken together, these analyses support predictions #1 and #2 above. Moreover, the confirmation of these predictions from the activity of relatively small numbers of neurons (387 total neurons in Fig. 2) shows that the grid cell population performs this decorrelation efficiently, thus supporting prediction #4 above.

Prediction #3 (controllable kernel width) is a natural consequence of the well-known increase of grid cell firing field widths when moving from dorsal to ventral rodent mEC. Probing this prediction more quantitatively requires averaging together sets of neurons based on their anatomical position along the dorsal-ventral mEC axis. To examine this prediction in more detail, we used recent population recordings from the study of Campell et al. (2021)^42^. This study contains recordings from the medial entorhinal cortex of mice traversing a one-dimensional virtual reality track along with anatomical locations of the neurons along the dorsal-ventral axis. Thus these data allowed us to analyze the effect of reading out from an anatomically-constrained subset of the neurons.

Grid cells can be hard to identify in 1D, and thus the neurons we analyze from this study are putative grid cells—they were identified in the study based on their tuning to distance run in the dark, and show a number of properties matching grid cells. Moreover, analyses of a separate dataset by the same authors revealed that such distance-tuned data are indeed grid cells^42^.

Following Campell et al. we analyzed these neurons in the dark to focus on pure encoding of internally-estimated position without the confounding effect of visual landmark input, and investigated their autocorrelation over short distances to minimize the effect of population drift in the dark. Campell et al. find that these neurons show periodic autocorrelations, with the period of the autocorrelation increasing along the dorsal-ventral axis, as expected for grid cell modules. For completeness, we regenerated the autocorrelation plots from this study, shown in Fig. 3a.

**Figure 3.**
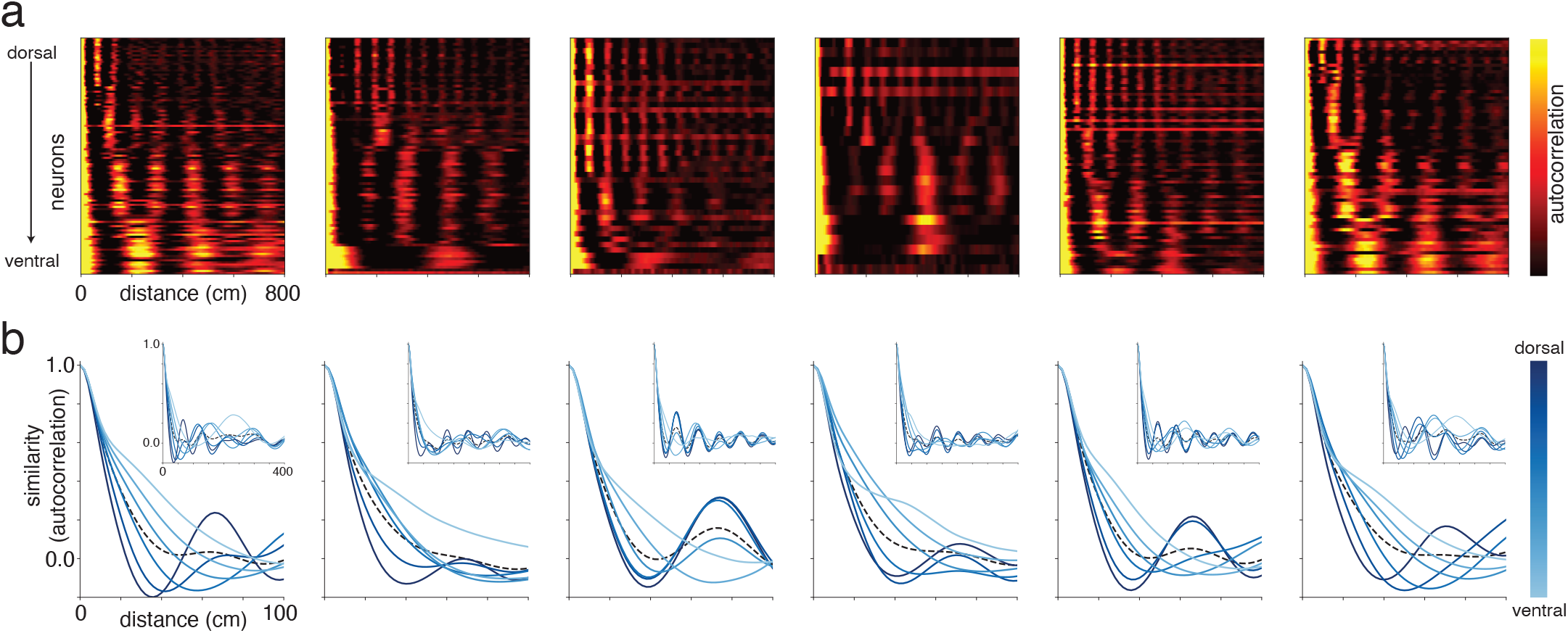
Similarity structure of putative grid cell representation from study of Campbell et al. (2021). (a) Autocorrelation of distance-tuned neurons for 6 recording sessions (left to right), showing periodic autocorrelation structure and a gradient of firing field widths and period along the dorsal-ventral axis. Note that the first 4 plots are the same plots shown in Fig. 6E of Campbell et al. (2021), regenerated from the data shared by the authors (and we choose similar formatting for consistency). (b) Similarity kernel of the grid cell representation (measured by autocorrelation) against physical distance between pairs of locations. Each sub-panel shows a different recording session (corresponding to sessions in (a)). Each trace within a sub-panel shows a subset of neurons grouped by anatomical position along the dorsal-ventral axis, with darkest blue showing most dorsal neurons and lightest blue showing most ventral neurons. Inset shows similarity kernel over longer distances. Note similarity to Fig. 1f.

To capture the effect of averaging together an anatomically-constrained subset of grid cell modules, we considered subsets of these neurons grouped by their position along the dorsal-ventral axis. Each subset contains 25% of the neurons in the population, and we considered 6 such overlapping subsets (0-25th percentile, 15-40th percentile, and so on). We computed the similarity of the putative grid cell representation as the averaged autocorrelation function for neurons within each subset, and plot these similarities as a function of distance in Fig. 3b. As can be seen from the plots, the similarity kernel shows a gradient of widths along the dorsal-ventral axis, suggesting that the threshold for similarity preservation vs. decorrelation can be easily controlled by anatomically-localized connectivity, as in prediction #3 above.

To summarize, these analyses of entorhinal cortex population recordings confirm that the similarity structure of grid cell responses acts to scramble physical distances.

### 2.4 Pattern separation in higher-dimensional and abstract spaces

Periodic, grid cell-like responses have been found not just while navigating physical space, but when performing tasks that involve navigating nonspatial sensory and abstract domains, suggesting a more general role in neural computation^4;8;44;45^. We note that similarity kernel estimation and pattern separation in two-dimensional abstract spaces function much as in two-dimensional physical space, and thus the conclusions from the previous sections hold in abstract 2D spaces.

To demonstrate how grid cell pattern separation might be used in abstract spaces of dimension higher than 2, we consider an architecture proposed by Klukas et al.^46^. This architecture suggests that two-dimensional grid cell modules can be used to encode a higher-dimensional space by first randomly projecting the space onto a set of planes and then using standard grid cell coding in each plane. That is, given a *D*-dimensional input space along with *M* grid cell modules, first construct *M* different random projections from *D* dimensions to 2 dimensions. The *m*th module then encodes the two-dimensional output of the *m*th projection as if it were regular 2D space. Each module thus forms a representation of the higher-dimensional space that is ambiguous both because of the projection into two-dimensional space (multiple points in the original space are projected onto the same point in the two-dimensional space) and because of the periodicity of the grid cell response (multiple points in the two-dimensional space cause the same grid cell population response). However, in general the existence of multiple modules guarantees that large high-dimensional spaces can still be uniquely represented by the grid cell population.

Much as in the 2D case, the grid cell representation of these higher-dimensional spaces also acts to scramble distances, meaning that except for very nearby points, distances in grid cell space do not reflect distances in the original space.

To concretely illustrate this pattern separation of an abstract high-dimensional space, we use as an example the latent space of a generative model trained to generate artificial faces (specifically, PGAN^43^, a type of Generative Adversarial Network^47^). The latent space of this model learns an abstract underlying representation of faces. Thus, each point in the latent space corresponds to a particular face. The latent space is moreover smooth and structured—nearby points correspond to similar faces, and a continuous trajectory between two points in this latent space interpolates between the faces corresponding to points at either end of the trajectory, Fig. 4a,b.

**Figure 4.**
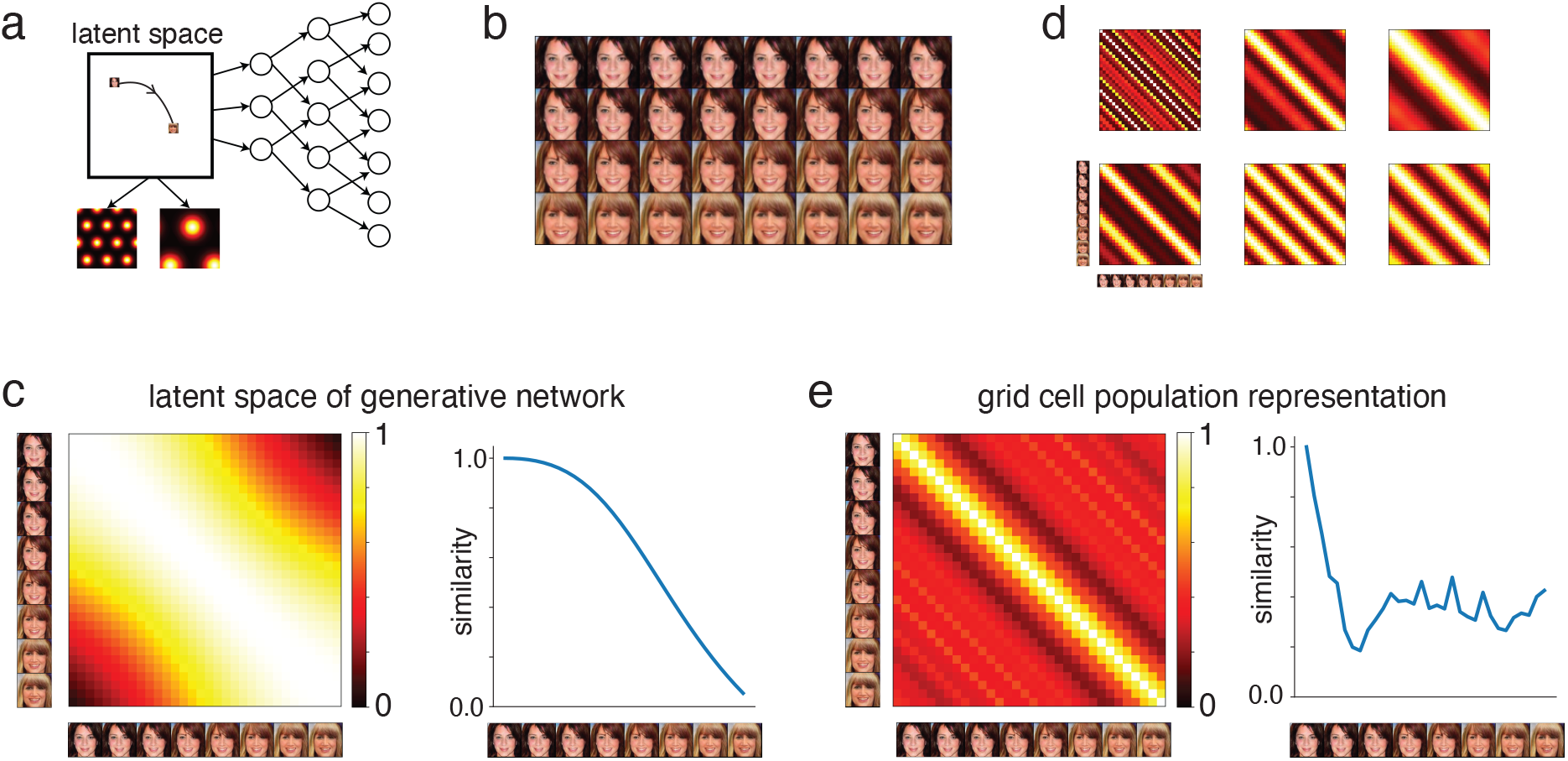
Grid cell pattern separation of latent representations in a generative model. (a) Schematic of a Generative Adversarial Network (PGAN^43^) that maps vectors from a latent space into image space to generate faces. The representations in the latent space are smoothly structured, with nearby points corresponding to similar faces. These latent representations are also passed through a grid cell layer (shown below the latent space) to generate representations analyzed in panels d and e. The smooth trajectory in the latent space illustrates the trajectory used to generate faces in panel (b). (b) Faces generated by a smooth trajectory through the latent space of the GAN. Trajectory moves from top left image to bottom right image. (c) Similarity kernel for pairs of latent vectors along the trajectory. Left: similarity for all pairs of latent vectors. Right: Similarity of first latent vector (i.e., first image) to remaining latent vectors in the trajectory. (d, e) Similarity kernel of grid cell representations, using a 6 module network. (d) Each plot shows similarity kernel constructed from a single module. Note that while the representations of nearby images are separated (i.e., dark bands immediately off the main diagonal, corresponding to small values), some distant images have overlapping representations (i.e., bright bands further off diagonal, corresponding to large values). (e) Similarity kernel of full grid cell population vectors (i.e., across all modules) corresponding to images. Left: similarity for all pairs of grid cell population vectors across trajectory. Right: Similarity of first grid cell population vector (i.e., first image) to remaining population vectors in the trajectory. Note the sharply peaked similarity, indicating decorrelation, as well as the lack of overlapping representations elsewhere on the trajectory (no bright off diagonal bands).

We pass the population activity vectors corresponding to points in this latent space as inputs to simulated grid cell populations. Despite the smoothly varying and highly similar structure of the input trajectory, we show that the resulting grid cell population representations are well-separated, with a similarity kernel peaked about the origin and then sharply decaying to a near-constant baseline, Fig. 4c,d.

Thus, even for this abstract high-dimensional space, grid cell responses are well-suited to decorrelate input patterns and scramble similarity structure.

## 3 Discussion

Theoretical analyses of the properties of the grid cell population code have highlighted a number of remarkable properties, among which is their ability to efficiently decorrelate or pattern separate the representation of spatial positions, except on very local scales^11;29;34^. We showed that this decorrelation can be captured by the use of a kernel function, *k*, that encodes the similarity of the encoding of pairs of points. Decorrelation implies that pairwise distance relationships are scrambled and homogenized, so that, except for very nearby points, *k*(***x***_**1**_, ***x***_**2**_) is small and bears no obvious relationship to the distance between ***x***_**1**_ and ***x***_**2**_. That is, there exist nearby locations with well-separated population vectors and distant locations with close population vectors, and a given value of population vector similarity (*k*) corresponds to a disparate set of spatial distances. Using neural population recordings from mouse and rat entorhinal cortex, we empirically confirmed the theoretically predicted structure of the similarity kernel, thus supporting a role for grid cells in decorrelating spatial representations while preserving very local structure. We moreover showed that such a pattern separation argument applies to proposed higher-dimensional grid cell architectures.

### Caveats and assumptions

It is important to emphasize that, in the absence of noise and distortions to the periodicity of grid cell firing, the grid cell representation does contain information about global distances. Thus it is possible to calculate the distance between two arbitrary points ***x***_**1**_ and ***x***_**2**_ from the grid cell population activity corresponding to these points (up to some large distance range) as well as to plan navigational routes using these representations ^11;15–19^. However, these distances are encoded in a hard to access, highly nonlinear way, and calculating long-range distances and navigational routes requires exploiting higher-order structure in the grid cell population representation. Thus our argument against the use of the grid cell representation in such global navigation relies on the difficulty and noise-sensitivity of such a computation when carried out by networks of biological neurons (rather than its impossibility), requiring large numbers of neurons, very large weights, or complex decoding strategies.

We make use of a particular notion of similarity between neural population activity (i.e., the function *k*(***x***_**1**_, ***x***_**2**_)), measured by the dot product between population vectors. Our results do not depend on specific quantitative features of the relationship of *k* with physical distance, but on qualitative features such as its non-monotonicity and the narrow spread of values of *k* for points that are not too close. While the exact shape of the non-monotonicity, the threshold distance, and the specific spread of similarities will change for different measures of similarity, the qualitative conclusions will hold for a wide range of similarity measures. In particular, in finite dimensions all ways of measuring the length of a vector are equivalent up to a scaling constant.^b^ Consequently, when switching to these alternative distance measures very nearby points still tend to remain close and distant points tend to remain far. Our results thus generalize to the wide class of similarity measures that in some way measure the length of the difference in population activity vectors.

### Role of grid cells in local navigation

Recent work has highlighted distortions of the regular hexagonal grid pattern, and used these distortions to argue that grid cells may play a role in local rather than global navigation^26^. Our observations that the multiperiodic grid cell representation is not well-structured to read out long-range metric and vector information, even in the absence of distortions of the grid, provide complementary support for such a role in forming very local maps. A local map can provide two important readouts for navigation. First, it can provide a unique distributed representation for different locations, which supports recognition of specific locations and encoding of other information associated with those locations. Second, it can provide local metric information near a given location, that supports local navigation to orient and navigate over relatively short distances when near that location (or imagining being near the location). By contrast, a global map puts much stronger constraints on the underlying representations, which need to provide self-consistent, stable and easily extractable metric information over a much broader scale.

### Local maps, global pattern separation, and episodic memory

A local map and a global scrambling function align well with the hippocampus’ role in episodic memory: encoding the specific details of individual experiences. In contrast, the relationships among distinct episodes—such as their relative temporal ordering—are often much harder to remember. Each episode can thus viewed as a local map, in which the relationships among the elements present in that episode are clearly specified. What is lacking, however, is a comprehensive global map that situates every episode in relation to the others. Establishing such cross-episode relationships requires explicit integrative processes that link details across different memories.

### Locality-preserving hash functions instead of coordinate systems as a conceptual metaphor for grid cells

The lattice structure of grid cell responses has led to suggestions that they act like a coordinate system, providing a global metric for space and enabling the computation of spatial displacement vectors and angles for navigation^1;9–15^. However, decorrelation by the grid cell population means that the “coordinates” assigned to nearby points are very different from each other, making this coordinate system poorly suited to computing distances between well-separated points and planning long-range navigational routes. Observations that grid cells form an efficient and highly decorrelated representation of a large input space suggest the alternative metaphor of hash functions in computer science^34^. Hash functions provide scrambled low-dimensional representations of high-dimensional input, assigning very different labels to nearby points. Much like grid cells, the mapping is many-to-one and the high capacity comes at the cost of occasional “collisions”, where a hash function maps very different inputs to the same output (the equivalent of collisions for grid cells is multiple well-separated spatial locations being mapped to the same grid cell population activity, as a consequence of the multiperiodic structure). Again, much like grid cells, these collisions are rare and usually not of practical concern. More generally, hash function-like computations have long been suggested as a key function of the hippocampal formation, inspired by its small size and role in rapidly learning conjunctive representations^48–50^.

While standard hash functions aim to decorrelate any pair of inputs, the grid cell population code preserves very local similarity structures—where “very local” refers to the scale of a single neuron’s firing field. The preservation of very local structure is evident in the smooth decay of the similarity kernel at short length scales (e.g., around 10–20 cm in the plots of Fig. 2c). This preservation suggests that the grid cell computation could in fact be an example of variants of hash functions called *locality-sensitive hash* (LSH) or *locality-preserving hash* (LPH) functions^34^. Both LSHs and LPHs decorrelate inputs that differ beyond a certain threshold, but LSHs map highly similar inputs to the same output (effectively carrying out a “same or different” computation with a threshold for similarity) while LPHs map similar inputs to nearby outputs, preserving local distance relationships.

The modular structure of grid cells closely resembles common strategies used in locality-sensitive hashing (LSH). Many LSH algorithms rely on a set of functions that map similar inputs to similar outputs while decorrelating most input pairs. However, these functions can sometimes fail by mapping different inputs to the same output. Crucially, such failures typically occur on different inputs for each function, so when combined, the set as a whole forms an effective LSH. These functions are thus analogous to the modules in the grid cell representation, where each module functions like a weak LSH, while their combination forms a strong LSH. Intriguingly, the grid cell representation is a novel LSH in that the definition of locality varies systematically across the component functions due to the multiscale nature of the modules. Furthermore, environmental features and rewards can distort the grid, suggesting that this hash function may be adaptive—potentially enhancing resolution or pattern separation in specific contexts. The grid cell code may thus be a useful source of ideas for designing adaptive or domain-specific locality-sensitive hash functions.

Locality-preserving hashing behavior, on the other hand, is compatible with the local map picture described above and might allow the grid cell system to provide a coordinate system and metric for very local spatial patches^26^. In this view, grid cells serve as coordinate systems on very local scales and as hash functions on intermediate to larger scales. The transition between these regimes would vary systematically by anatomical location, depending on the width of grid cell firing fields. Thus, locality-preserving hashes might provide a useful conceptual metaphor for grid cells intermediate between global coordinate systems and pure hashing, allowing a reconciliation between these perspectives.

## 4 Methods

### 4.1 Grid cell models

We consider phenomenological tuning curve models for the grid cells, meaning that the response of each neuron is given by some tuning curve function *s*(***x***), where *s* is a periodic function of the position ***x***. Such tuning to position could emerge from sensory input corresponding to landmarks combined with integration of a self-motion velocity signal, and could be maintained by recurrent attractor dynamics. However the model seeks to describe grid cell responses rather than account for their mechanistic origin. Neurons are grouped into a small number of modules, each distinguished by a characteristic period and orientation for the tuning curve function, but with different phases. The different models are distinguished by the shapes of the tuning curve functions.

### 1D grid cell model

Each module in the 1D grid cell population is defined by a characteristic scalar period, *λ*_*i*_. The response of the *j*th neuron in the *i*th module at position *x* is given by a von Mises tuning curve as 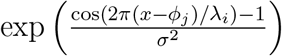, where *ϕ*_*j*_ is the neuron’s phase and *σ* is the field width, chosen to be 0.55. For the plots in Fig. 1a we show two modules with periods 0.7 and 1 respectively and use 50 neurons per module. Each entry of the correlation matrix is the mean-subtracted correlation between the population vectors at two points. To generate the kernels for each module we plot the normalized dot products ***g***(0) ***g***(*x*)*/*(||***g***(0) ||_2_ ||***g***(***x***) ||_2_) for different values of *x*. The plots in Fig. 1b are generated similarly except we construct population vectors using 8 modules with periods [0.46, 0.7, 0.87, 1., 1.3, 1.7, 2.21, 2.87].

### 2D grid cell model

The 2D grid cell model is based on the model used by Klukas & Fiete (2020). For each module, a unit hexagonal lattice Λ is defined using the two basis vectors [1, 0] and [cos(*π/*3), sin(*π/*3)]. The phase of a particular spatial position ***x*** is defined to be ***ϕ***_***x***_ = *x/λ*_*m*_ mod Λ, where *λ*_*m*_ is the period of the module. Tuning centers for the neurons are chosen to uniformly cover a cell of this lattice. The response of a neuron with preferred phase ***ϕ*** to position phase ***ϕ***_***x***_ is Gaussian of the form *A* exp (−*d*(***ϕ, ϕ***_***x***_)*/σ*^2^), where *d* is the distance computed modulo the lattice, *A* is the maximum firing rate and *σ* is the field width. For simulations we chose *A* = 20 and *σ* = 0.15. For the plots in Fig. 1c we show two modules with periods 1 and 2.21 respectively and use 16 neurons per module. We calculate firing rates on a 200 × 200 grid evenly spaced in a 3 × 3m arena. Kernels are calculated as for the 1D case. For panel *d* we use 8 modules of periods [0.46, 0.6, 0.77, 1., 1.3, 1.7, 2.21, 2.87]. For panel *e* we first randomly shuffle the modules (yielding the order [4, 0, 6, 2, 5, 7, 3, 1], where the modules are numbered 0 through 7). We then plot the average kernel for the first module, the combined first two modules and all modules respectively. For panel f, we plot the average kernels for modules 0 through 4 (shorter period) and modules 4 through 8 (longer period).

### Higher-dimensional grid models

The higher-dimensional grid cell model is also based on the model used by Klukas & Fiete (2020). It is identical to the 2D model except for an initial random projection step from the *M*-dimensional input space to a 2D plane. This is implemented as a set of 2 *× M* random matrices *P*_*m*_, one for each module. Each matrix has IID zero mean Gaussian entries with standard deviation 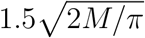 This choice of standard deviation approximately preserves the length of vectors, so that all scaling comes from the grid cell period.

### 4.2 Analysis of Gardner et al. (2022) data

The data from Gardner et al. contains recordings from 2 rats that have more than 1 module: 2 days of recordings from 3 modules of rat R and 1 day of recording from 2 modules of rat Q. We used functions from the code accompanying these data to load the positions and spike times for grid neurons from the various modules.

We then computed the mean firing rate for each neuron in 20 cm by 20 cm spatial bins covering the recording arena, yielding a 2D tuning curve for each neuron. The kernel *k*^*m*^(*x*_1_, *x*_2_) for two spatial locations *x*_1_ and *x*_2_ and module (or combination of modules) *m* is then calculated as ***g***^***m***^(***x***_**1**_) · ***g***^***m***^(***x***_**2**_)*/*(||***g***^***m***^(***x***_**1**_)||_2_||***g***^***m***^(***x***_**2**_) ||_2_), where the *n*th entry of ***g***^***m***^(***x***_**1**_) is the tuning curve of the *n*th neuron at spatial location ***x***_**1**_. Note that ***x***_**1**_ and ***x***_**2**_ are 2D locations and thus this kernel can also be interpreted as a four argument function *k*^*m*^(*x*_1_, *y*_1_, *x*_2_, *y*_2_), where ***x***_**1**_ = (*x*_1_, *y*_1_) and ***x***_**2**_ = (*x*_2_, *y*_2_).

The values of these kernels for specific locations are shown in Fig. 2a,b. That is, we plotted *k*(***x***_**0**_, ***x***), where ***x***_**0**_ is the center of some chosen spatial bin and ***x*** ranges over all the spatial bins in the arena.

For plots in Fig. 2c, we considered all pairs of spatial bins, centered at ***x*** and ***y*** respectively, and their accompanying kernel values *k*^*m*^(***x, y***). We grouped pairs of bins by the Euclidean distance between their centers (i.e., ||***x*** − ***y***||_2_) and plotted percentiles of the distribution of kernel values in each group. For plots in Fig. 2d, we plotted the distributions of kernel values for all pairs of points between 0.3 and 2m apart, gathered into 0.1m bins.

For the plots in Fig. 2e, we consider a separate threshold linear classifier (perceptron) for each position ***x***_**0**_. Given a threshold *θ* and an input population vector ***v***, the classifier reports that the animal is at position ***x***_**0**_ when 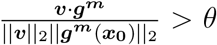. We applied this decoder to the mean firing rates for each position bin ***x, g***^***m***^(***x***). The decoding error for position ***x***_**0**_ was then computed as the average Euclidean distance from ***x***_**0**_ of all points ***x*** for which the decoder reports that the animal is at ***x***_**0**_. We then repeated this process to yield a decoding error for each location. Note that this decoder is equivalent to applying the threshold to *k*^*m*^(***x***_**0**_, ***x***).

### 4.3 Analysis of Campbell et al. (2021) data

Fig. 3a is a reproduction of results shown in Fig. 6E from Campbell et al. (2021), which we regenerated using scripts publicly shared by the authors. The original figure showed 4 sessions from the data, and we added 2 more sessions for a total of 6 sessions. To generate Fig. 3b, we grouped neurons by anatomical location using a sliding window on depth. Each group contains 25% of the neurons and each successive group shifts the start location by 15% (i.e., first group contains 0-25th percentile of depth, second group contains 15-40th percentile of depth, and so on).

### 4.4 Pattern separation of higher-dimensional abstract spaces

For the generative model plots in Fig. 4 we used the PGAN architecture of Karras et al. (2017)^43^ and downloaded pretrained model weights from Pytorch Hub^51^. We augment this model with a set of 7 grid cell modules, modeled using the architecture of Klukas & Fiete (2020) (described under the section of grid cell models). Activity from the latent space is randomly projected into each of these modules, using a separate random projection. The modules have periods of [1., 2.1, 3., 3.8, 4.2, 5.3, 6.1], where units are chosen so that the typical separation between random vectors in the original latent space is 10 units.

Fig. 4b shows the generative model output corresponding to a smooth one-dimensional trajectory through the latent space of this model. For the plots in Fig. 4c-e we calculated the kernels as the normalized dot products of the representations in the appropriate layer of the model, just as for the kernels in the other figures.

## Acknowledgements

We thank Ila Fiete and Christopher Kymn for helpful discussions. This material is based upon work supported by the Air Force Office of Scientific Research (AFOSR) under award number FA9550-22-1-0532 (R.C. and R. O.) and by the Navy Office of Naval Research under award number N000142412325 (C.R. and R.C.).

## Footnotes

a Note that the notation ***g***(***x***) simply indicates that the population activity depends on ***x*** and does not indicate that the input to the grid cell population is itself spatial location. The actual input could be a combination of landmark input and a velocity signal that is integrated to update the grid cell representation from ***g***(***x***_**1**_) to ***g***(***x***_**2**_) as the subject moves from ***x***_**1**_ to ***x***_**2**_

b More technically, all norms impose the same topology, and in high dimensions concentration of measure tends to linearize relationships between norms

